# Genotyping sequence-resolved copy number variation using pangenomes reveals paralog-specific global diversity and expression divergence of duplicated genes

**DOI:** 10.1101/2024.08.11.607269

**Authors:** Walfred Ma, Mark JP Chaisson

## Abstract

Copy number variant (CNV) genes are important in evolution and disease, yet sequence variation in CNV genes remains a blind spot in large-scale studies. We present ctyper, a method that leverages pangenomes to produce allele-specific copy numbers with locally phased variants from next-generation sequencing (NGS) reads. Benchmarking on 3,351 CNV genes, including *HLA*, *SMN*, and *CYP2D6*, and 212 challenging medically relevant (CMR) genes that are poorly mapped by NGS, ctyper captures 96.5% of phased variants with ≥99.1% correctness of copy number on CNV genes and 94.8% of phased variants on CMR genes. Applying alignment-free algorithms, ctyper requires 1.5 hours per genome on a single CPU. The results largely improve predictions of gene expression compared to known expression quantitative trait loci (eQTL) variants. Allele-specific expression quantified divergent expression on 7.94% of paralogs and tissue-specific biases on 4.68% of paralogs. We found reduced expression of *SMN2* due to *SMN1* conversion, potentially affecting spinal muscular atrophy, and increased expression of translocated duplications of *AMY2B*. Overall, ctyper enables biobank-scale genotyping of CNV and CMR genes.

## Introduction

Human genomes are characterized by frequent duplications and deletions, leading to copy number variation. Up to 10% of protein-coding genes are known to be copy-number variable, showing distinct distributions across human populations^1,2^ and association with traits such as body mass index^3^ and disease, including cancer^4^, cardiovascular diseases^5^, and neurodevelopmental disorders^6,7^. While copy number variants (CNVs) are infrequent genome-wide, regions of long, low-copy repeats called segmental duplications (SDs) are enriched in genes and are catalysts for recurrent CNVs due to non-allelic homologous recombination^8–10^. These regions include *TBC1D3*, *NPIP*, and *NBPF*^11,12^ with particular impact in humans through association with brain function and adaptation^13–16^. SD regions not only undergo frequent copy number changes but also experience more rapid structural alterations and exhibit an elevated rate of nucleotide substitutions compared to nonrepetitive DNA^17^. As a result, studies that only consider the aggregate copy number (aggreCN) of duplicated genes miss non-reference gene duplicates and variation among multiallelic CNVs, which have both been shown to influence phenotypes and disease susceptibility^18–20^, including hypertension and type 2 diabetes^21^.

There are limited studies on sequence variation between gene duplicates, particularly in studies using short-read next-generation sequencing data. Existing CNV-calling tools detect excess coverage on reference alignments, hiding variants among copies^22^. Furthermore, alignments to a single reference contain ambiguity and bias^23^ and are unable to detect novel duplicates. Advances in single-molecule sequencing enabled the assembly of pangenomes from diverse populations^24–26^. Almost all novel sequences found in the pangenome are CNV events^24^, providing sequence-resolved CNVs and non-reference duplications. While graph-based pangenomes reduce reference bias^27^, they merge similar sequences—including alternative alleles and paralogs from distinct genomic locations and functions—into shared graph paths, obscuring paralog-specific variants and novel duplications^28^. Furthermore, as pangenomes include more samples, the alternative sequences, gene conversion, and genome rearrangements that are prevalent in divergent genomic regions present an even greater challenge to be represented as graphs^17^, motivating the need to develop tools to analyze duplicated genes using NGS reads with pangenomes.

Here, we present ctyper, a method that directly compares NGS reads to unmerged pangenome haplotypes to identify the most similar genomic segments corresponding to genes in an NGS sample and determine their copy number states. By avoiding sequence merging, ctyper preserves locally phased variants and captures complex variation, such as structural variation and gene conversion, that are often missed in NGS analyses. In this study, we focus on sequences of complex, duplicated genes that would otherwise be challenging to analyze using NGS data and canonical reference genomes. Leveraging alignment-free techniques, and introducing a novel mathematical model for genotyping haplotypes with a polynomial-time solution, ctyper achieves both high accuracy and computational efficiency, making it well-suited for large-scale biobank data analyses.

## Results

### Overview of the genotyping method

Our method, ctyper, genotypes sequence-resolved copy number variation in NGS data leveraging haplotype-resolved assemblies. Instead of representing genetic variation in the form of individual variants, we represent variation as haplotype segments that are short enough to minimize disruption by recombination, allowing precise sharing with an NGS sample through identity-by-descent^29^, and long enough to capture structural information of genomic sequences. We refer to these haplotype segments as pangenome-derived alleles (PAs). PAs capture phased variation, representing structural variation, gene conversion events, and small variants. In this study, we set the boundaries of PAs to study the CNVs of protein-coding genes. Each PA includes consecutive exons separated by <20 kb, with an additional 5 kb of upstream and downstream sequences added as anchors (Methods). The 20 kb cutoff reflects both functional proximity, as most short-range transcription factors operate within this range^30^, as well as population-level genomic linkage. PAs typically range from 10 to 100 kb, corresponding to the scale of linkage disequilibrium (LD) blocks^31^, within which most single-nucleotide polymorphisms (SNPs) are in strong LD and can be treated as single units of inheritance. While PAs generally correspond to individual genes, they also may cover only a fraction of a gene flanked by long introns or, conversely, include paralogs within the 20 kb boundary.

For computational efficiency and to avoid alignment ambiguity in repetitive DNA, we use an alignment-free comparison of low-copy *k*-mers (DNA fragments of a fixed length *k*, here *k* = 31) measured in query NGS samples to genotype PAs. For each gene of interest, we group all similar PAs in the pangenome, including orthologous genes, paralogs, and homologous pseudogenes, to construct a matrix to be used in genotyping that contains the *k*-mer composition of all similar PAs (Methods). The rows of a matrix correspond to individual PAs, while columns correspond to *k*-mers found within the group of PAs and not elsewhere in the genome. The values of the matrix reflect the multiplicity of a *k*-mer in each PA (Methods; Figs. 1a,b).

**Figure 1.**
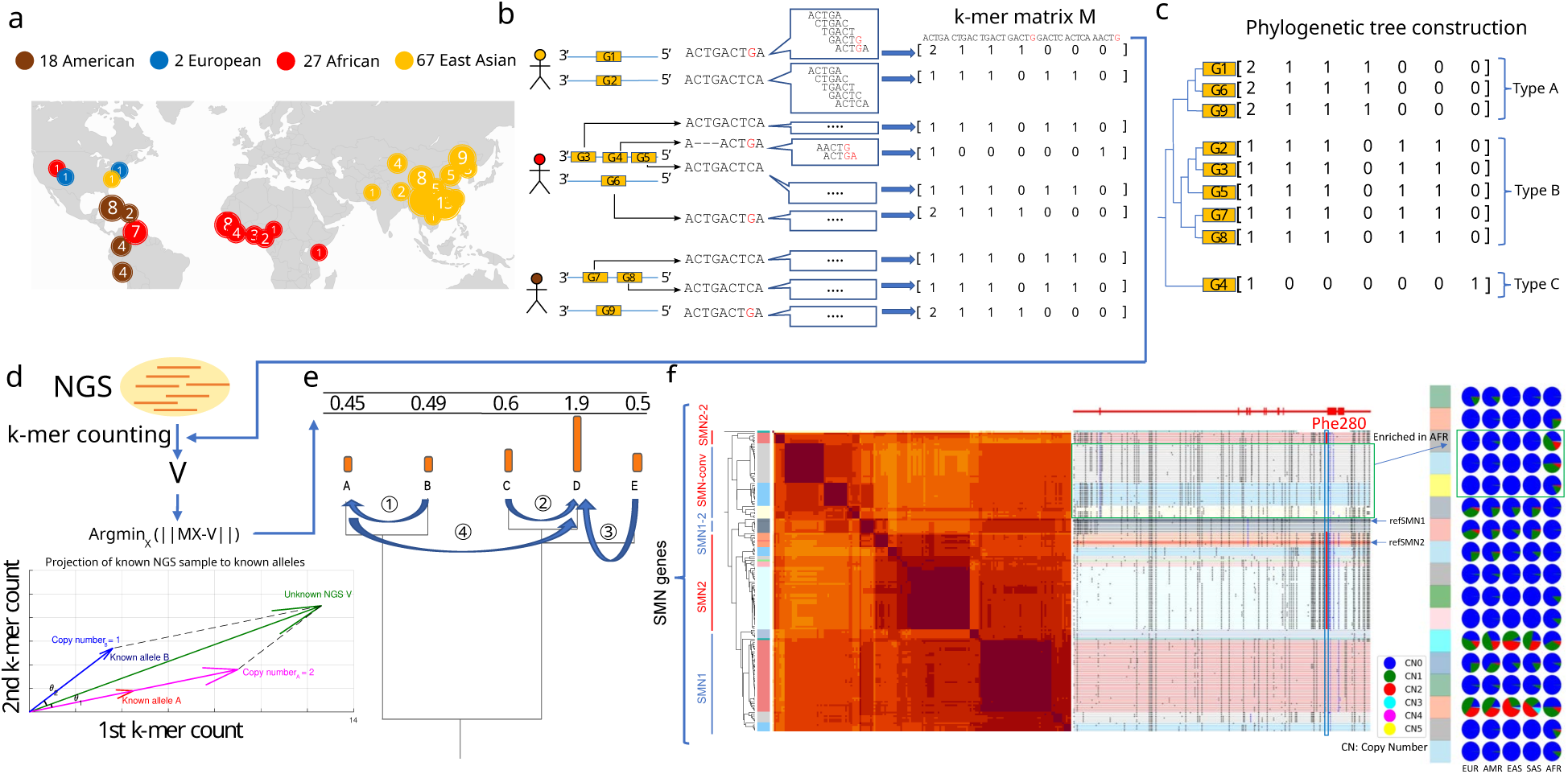
**a**, Demography of the reference pangenome assemblies. **b,** Construction of pangenome *k*-mer matrices for CNV genes. Each individual gene is represented as a vector of counts of *k*-mers exclusively found among homologous sequences. All similar sequences including paralogs and orthologs are included and integrated as a *k*-mer matrix. **c,** Construction of phylogenetic trees based on *k*-mer matrices. **d,** Schematic of approach to estimate genotypes of alleles using NGS data. The *k*-mers from each matrix are counted in NGS data and normalized by sequencing depth. The normalized *k*-mer counts are projected to all pangenome genes. **e,** Reprojection of the raw results in the last step to integer solutions recursively based on the phylogenetic tree. **f,** An illustrative annotation and genotyping results on *SMN1/2* genes using HPRC samples. All *SMN* genes are categorized into five major types and 17 subgroups. *SMN1*/*SMN2* correspond to the most common types of each paralog; *SMN1-2*: *SMN1* partially converted to *SMN2*; *SMN*-conv: additional converted SMN genes *SMN2-2*: a rare outgroup of *SMN2*. *SMN*-conv is predominantly mapped to the *SMN2* locus and is found to be enriched in African populations. The GRCh38 assembly includes *SMN1-2* and *SMN2*. On the right side of the classification, the phylogenetic tree and heatmap of pairwise similarities are shown along with a mutant plot based on an MSA highlighting point differences to *SMN1* in CHM13. Phe-280, the variant found to disrupt the splicing of *SMN2* transcripts, is highlighted in red. The genotyping results of 1kGP continental populations are shown on the right. Rows correspond to subgroups, columns correspond to continental populations, and the colors of pie charts give distributions of copy numbers among each continental population.

The genotyping of an NGS sample is performed per matrix by identifying a combination of PAs (rows) and their allele-specific copy numbers such that their total corresponding *k*-mer counts are the least-squared distance to observed *k*-mer counts from the sample. To do so, the *k*-mer counts from a sample are projected into the vector space of each *k*-mer matrix, and then assigned integer copy numbers using recursive rounding based on the phylogenetic tree of PA sequences (Methods; Figs. 1c-e). The genotyping results are a list of PA-specific copy numbers (paCNs).

As an example, there are 178 PAs for *SMN* genes, the gene family associated with spinal muscular atrophy. This includes copies of *SMN1* and *SMN2* as well as paralogs that have undergone gene conversion^32^, for example paralogs mapped to the *SMN2* locus but containing the *SMN1* version of phe-280, the SNP responsible for dysfunctional exon 7 splicing of *SMN2*^33^ (Fig. 1f).

### Pangenome-derived allele database construction and annotation

We constructed a PA database for 3,351 genes previously reported as CNV^24,26^ (Supplementary Table 1), using 114 diploid PacBio-HiFi assemblies from the Human Pangenome Reference Consortium (HPRC), Human Genome Structural Variation Consortium (HGSVC), Chinese Pangenome Consortium (CPC), and two haplotype-phased telomere-to-telomere assemblies^34,35^, in addition to GRCh38 and CHM13^36^. PAs were extracted from assemblies and annotated with other similar PAs within the same matrices used for genotyping, for example, the amylase genes (Fig. 2a). In total, we defined 1,408,209 PAs and organized them into 3,307 matrices (Fig. 2b,c).

**Figure 2.**
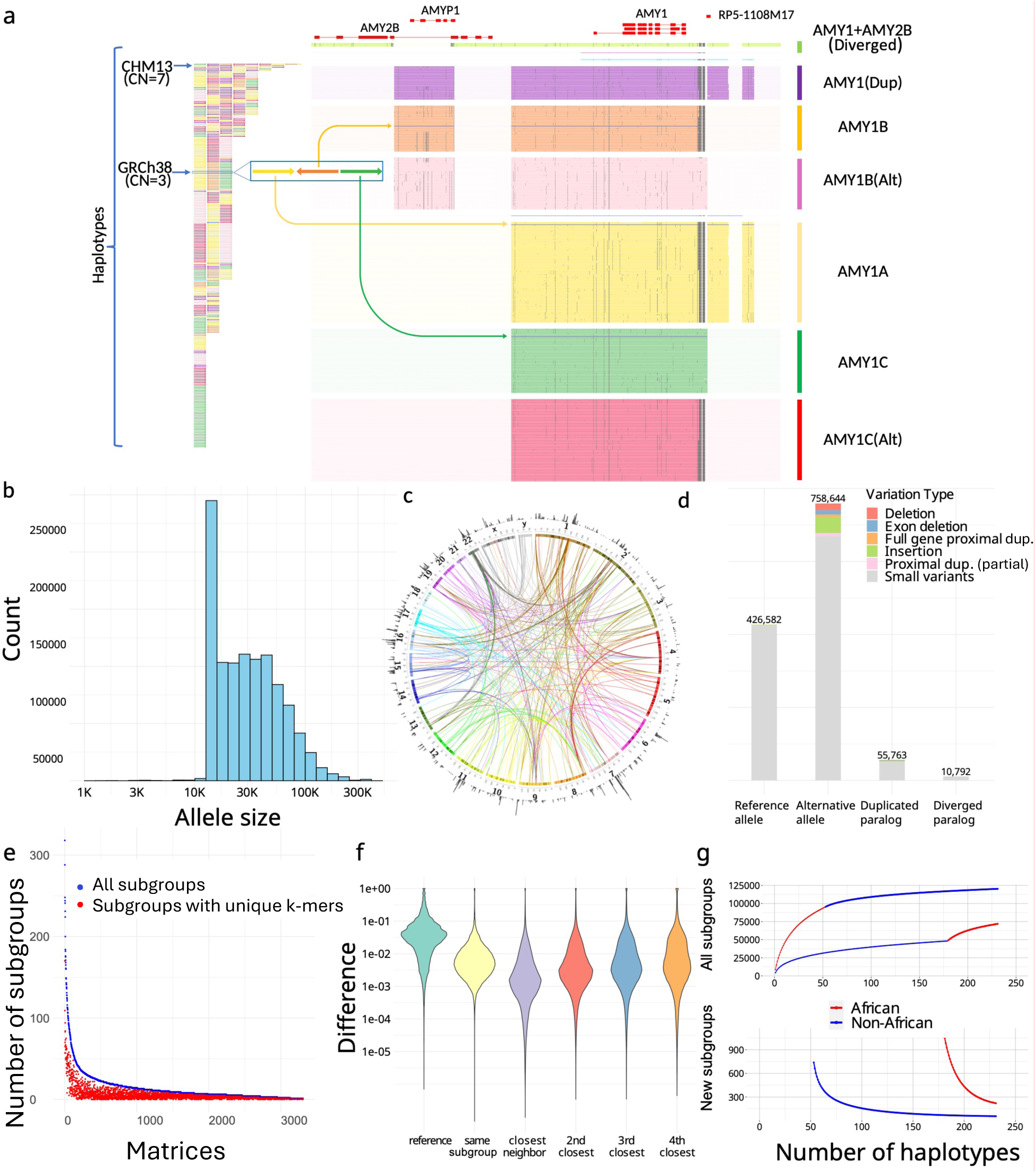
**a**, An overview of amylase 1 pangenome-derived alleles (PAs). (left) The corresponding order of all *AMY1* PAs on assemblies, which are colored based on their major groups (unions of subgroups devoid of SVs >300 bp between members). (right) *AMY1* genes are extracted as PAs as well as their flanking genes and sequences, including an *AMY2B* translocated proximally to *AMY1*, and two pseudogenes: *AMYP1* and *RP5-1108M17*. All PAs are vertically ordered according to the phylogenetic tree and aligned via graphic MSAs (Supplementary Methods). Homologous sequences are vertically aligned. Mutations are visualized as dots, and large gaps (deletions) are visualized as spaces. Seven major groups are categorized including five paralogs and two orthologs. There are no pseudogenes around *AMY1C*, while *AMY1A* has *RP5-1108M17* nearby and *AMY1B* has *AMYP1* nearby. There are alternative versions of *AMY1B and AMY1C*, with sequence substitutions. A new paralog called *AMY1(Dup)* is found primarily on haplotypes with duplications and has both pseudogenes nearby. Another paralog of *AMY1* found with translocated *AMY2B* is called *AMY1+AMY2B*. There are also two rare paralogs (blue and violet) and one singleton ortholog (steel-blue). **b,** The size distribution of PAs on a log density. The minimum size of PAs is 15 kb, though smaller alleles may be annotated on alternative haplotypes on GRCh38 and as partial loci when dividing large genes into alleles without recombination. **c**. CIRCOS plot of all PAs. (outer ring) The density of PAs in each Mb on GRCh38. (arcs) Interchromosomal PAs are included in the same group. **d,** Annotation of PAs according to orthology and variants with respect to GRCh38. **e,** Identifiability of highly similar subgroups by unique *k*-mers. The total number of subgroups (blue) and the number of subgroups that may be identified by paralog-specific *k*-mers (red) are shown for each matrix with a size of at least three. **f,** The distribution of logistic pairwise divergence of PAs depending on orthology and phylogenetic relationship. The values shown are average values from each of the matrices. Small neighbor distances are an indicator of the strong representativeness of the current cohort. **g,** Saturation analysis for all subgroups using a recapture mode according to two sorted orders: African genomes considered first, and non-African genomes considered first. The former order has a smoother curve than the latter order, indicating there are more African-specific subgroups.

Because the human population exhibits limited genetic diversity and LD is stronger within a short distance, PAs are often highly similar or identical. To reduce genotyping dimensionality and facilitate cohort analysis, particularly when sample sizes are limited, we used the phylogenetic relationship between sequences to merge similar PAs into highly similar subgroups (subgroups) to be treated as equal states (Methods). In total, we defined 89,236 such subgroups, which were used to enumerate all PA states observed in the pangenome, analogous to *HLA* nomenclature (Supplementary Figure 1).

To annotate low-frequency variants not represented by subgroups and annotate orthologous/paralogous relationships, we mapped PAs to GRCh38 genes (Supplementary Methods). The alignments provide sequence variation at a greater resolution than what is annotated by subgroups as well as the reference genomic locations. In total, 164,237 paralogous PAs across 6,389 loci were determined. The paralogous PAs that were similar to their corresponding reference locus (≥80% *k*-mer similarity) were labeled as duplicative paralogs, and the remaining lower-identity paralogous PAs were labeled as diverged paralogs. In total, 10,792 diverged paralogs from 2,734 subgroups were identified across 333 matrices (Fig. 2d). The divergent paralogs represent novel sequences recalcitrant to analysis using a single reference genome. For example, some amylase PAs include paralogs for both *AMY1* and *AMY2B* with low *k*-mer similarity to reference loci and were annotated as diverged paralogs, which reflects an *AMY2B* translocation event (Fig. 2a).

While most duplication events happen on different loci from their original genes, 6,673 PAs reflected proximal (<20 kb) duplications, including 1,646 PAs across 36 genes that exhibited “runway duplication”^37^ with at least three proximal duplications (Supplementary Figure 2). Proximally duplicated genes were grouped with their ortholog genes in the same PAs as they are unlikely to be inherited independently. Since ctyper can also genotype orthologous genes, we classified orthologous genes into two categories: reference alleles, which belong to the same subgroup as the reference gene, and alternative alleles, which fall outside of these subgroups (Fig. 2d).

### Pangenome-derived alleles and highly similar subgroups capture unique aspects of population diversity

Since other variant representations have been used in CNV studies, we assessed whether PAs capture unique aspects of genomic information that cannot be replicated by these methods. We compared paCNs to other CNV representations, including copy numbers of reference genes^1,37^, singly unique nucleotide *k*-mers^1,37,38^ (SUNKs), and large haplotype structures^39–42^. We found that PAs provide higher resolution of variation (e.g., SNVs), as 94.7% of variants are not reflected by the sequences of genes in GRCh38. Additionally, both nearby SUNK markers (Fig. 2e) and large haplotype structures were found to be poor proxies for PAs, and only a small proportion of paCNs were found to link to SUNKs or large haplotype structures (Methods).

We evaluated the extent to which global population diversity could be represented using subgroups compared to individual PAs. We found that even with largely reduced dimensions, subgroups still capture more than 80% of the total population variation (Fig. 2f). Additionally, we performed saturation analysis^43,44^ to estimate the percentage of the unobserved subgroups in each new genome. The current cohort represents 98.7% of subgroups in non-Africans and 94.9% in Africans (Methods), suggesting a near-saturated database (Fig. 2g).

### Genotyping pangenome-derived alleles among NGS samples and benchmarking results

We applied ctyper to genotype NGS samples from the 1000 Genomes Project (1kGP), including 2,504 unrelated individuals and 641 offspring. At the cohort level, excluding intronic/decoy PAs, bias was measured using Hardy-Weinberg equilibrium (Methods), and accuracy was measured using trio concordance (Methods; Supplementary Table 2). Overall, 0.75% of subgroups had significant disequilibrium (p < 0.05) (Fig. 3a) and 27 matrices having >15% subgroups with significant disequilibrium, which were mostly short genes (median = 4,564 bp) with fewer low-copy *k*-mers (Supplementary Table 3). The average F-1 score for trio concordance was 97.58% (Fig. 3b), while 18 matrices had high discordance (>15%), primarily for subtelomeric genes or on sex chromosomes (Supplementary Table 4).

**Figure 3.**
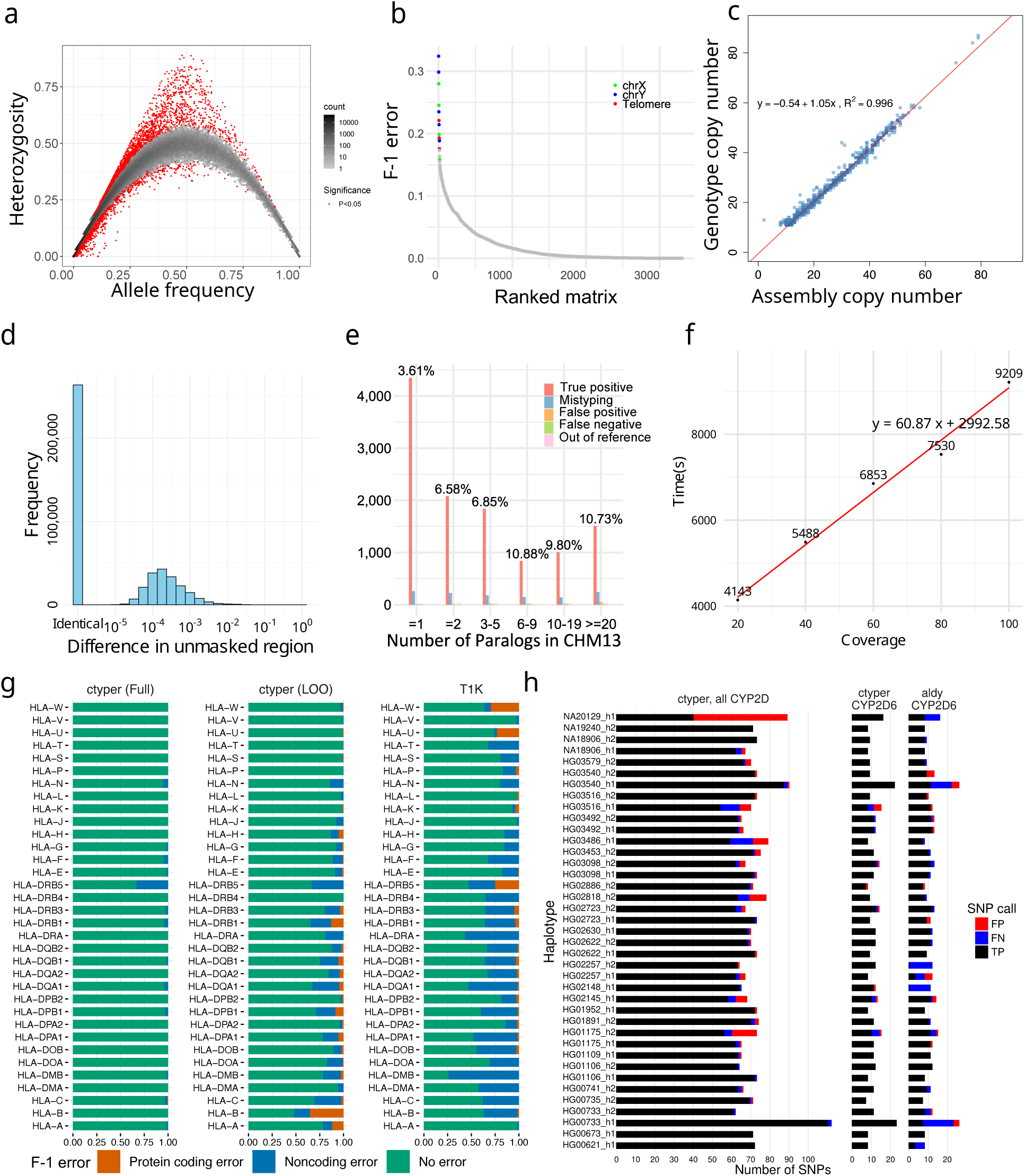
**a**, Hardy-Weinberg equilibrium of genotyping results on 2,504 1kGP unrelated samples. **b,** Genotype concordance of genotyping results on 641 1kGP trios, ordered by F-1 error. The matrices with an F-1 error of more than 15% were labeled by genomic location. **c,** Copy number comparison between assemblies and genotyping results on 39 HPRC samples shared with 1kGP. **d,** Sequence differences between genotyped and original alleles during leave-one-out test using pairwise alignment of nonrepetitive sequences. **e,** Detailed leave-one-out comparison in the diploid telomere-to-telomere genome CN1. The results are categorized regarding the number of paralogs in CHM13 to show performances on different levels of genome complexity and the main sources of errors. CN1 NGS sample was about 40 coverage. **f,** Runtime of ctyper on CN1 on all loci for varying coverage. **g,** Benchmarking of *HLA* genotyping using ctyper on full, and leave-one-out (LOO) databases, compared with T1K on 31 *HLA* genes. **h,** Benchmarking of *CYP2D* annotation on all *CYP2D* genes and *CYP2D6* exclusively.

We assessed copy number accuracy and bias in challenging and highly duplicated gene families, such as amylase, *NBPF*, *NOTCH*, *GOLGA*, and *TBC1D3*. The copy numbers derived from genotyping were compared to those in their corresponding assemblies for 39 HPRC samples shared with the 1kGP, using a database inclusive of these samples. For each sample, we benchmarked on all matrices where the corresponding assembly was high in copy number (>10 PAs) and no low-confidence sequences filtered during matrix construction (Methods). The results showed a high correlation (*ρ*=0.996, Pearson correlation) and little bias (Fig. 3c), with a 0.2% rate of missing copies (false negatives) and a 2.4% rate of additional copies (false positives), with a higher false positive rate likely explained by unassembled genes in the assemblies. When expanding benchmarking to include the remaining less challenging genes within matrices without filtered sequences on all 2,504 1kGP samples, the copy number between genotype and assemblies had an effectively identical correlation (*ρ*=0.996, Pearson correlation).

To evaluate sequence accuracy at the individual PA level, we performed assembly comparisons to assess the sequence similarity of the genotyped alleles to the sample assembly for the 39 HPRC benchmarking genomes. Each sample was genotyped with the full database (full-set) or the database excluding its corresponding PAs (leave-one-out). We assigned the genotyped PAs to the corresponding assembly (Methods), excluding introns/decoys and sequences with <1 kb non-repetitive bases, and measured the similarity between the genotyped allele and assigned query using global alignment^45,46^. We performed a similar analysis treating the closest neighbor to each assembly PA from the database as the correctly genotyped locus. Due to mismatching from database sampling or misassemblies, 2.9% of PAs from the leave-one-out and 1.0% from the full-set could not be paired to truth copies for assessment. Using the full-set database, paired PAs have 0.36 mismatches per 10 kb, with 93.0% having no mismatches on non-repetitive regions. The leave-one-out tests had 2.7 mismatches per 10 kb on non-repetitive regions, which was 1.2 additional mismatches per 10 kb from the optimal solutions (closest neighbors); 57.3% alleles had no mismatches, and 77.0% were mapped to the optimal solution (Fig. 3d). The leave-one-out results were 96.5% closer to the original PAs compared to the closest GRCh38 gene at 79.3 mismatches per 10 kb.

To isolate sources of errors in cases of misassembled duplications, we directly compared leave-one-out genotyping results to a telomere-to-telomere assembly^34^, filtering out intronic/decoy sequences. The sample genotypes had 11,627 correctly matched subgroups, 599 (4.8%) mistyped to other subgroups, 131 from singleton subgroups removed from the database during the test (1.1%; out of reference), 127 false positives (0.5% F-1), and 93 false negatives (0.4% F-1) for a total F-1 error of 6.7% (Methods; Fig. 3e). This is a 3% increase in mistypes compared to trio discordance. The copy number agreement was 99.1%.

The computational requirements are sufficient for biobank analysis. The average runtime for genotyping 3,351 genes at 30× coverage was 80.2 minutes (1.0 min/1× coverage for sample preprocessing, and 0.9 s/gene for genotyping) on a single core (Fig. 3f) using ∼20GB RAM, with support for parallel processing.

We compared the benchmarking of *HLA*, *KIR*, and *CYP2D6* to the locus-specific methods T1K^47^ and Aldy^48^. For 31 *HLA* genes, ctyper had an F1-score of 98.9% in predicting all four fields of *HLA* nomenclature^49,50^ against the full-set and 86.3% among the leave-one-out, while T1K had 70.8%. For protein-coding products (first two fields), ctyper reached 99.98% against the full-set (with 99.9% copy number F1-correctness) and 96.5% (with 99.5% copy number F1-correctness) among the leave-one-out, and T1K had 97.2% (Fig. 3g) (Supplementary Table 5-6). For 14 *KIRs*, ctyper reached 98.5% of predicting full fields against the full-set and 70.6% among leave-one-out, while T1K had 32.0% due to the limited database. For protein-coding products (first three digits), ctyper reached 99.2% against the full-set (with 99.9% copy number F1-correctness) and 88.8% among leave-one-out (with 99.2% copy number F1-correctness), while T1K had 79.6% (Supplementary Figure 3). Benchmarking *CYP2D6* star annotations based on assemblies^51^, ctyper reached 100.0% against the full-set and 83.2% among leave-one-out, compared to 80.0% using Aldy (Fig. 3h). There was perfect agreement of SNP variants ctyper against the full-set and 95.7% among leave-one-out, compared to 85.2% using Aldy.

We compared genotyping F1-scores on *HLA* to a contemporary method also based on pangenomes called Locityper^46^. Locityper performs slightly better at predicting all four fields, including noncoding variants (leave-one-out: Locityper 87.9% vs. ctyper 86.3%; full-set: Locityper 99.5% vs. ctyper 98.9%), while ctyper is slightly better on the first two fields of protein-coding products (leave-one-out: Locityper 94.0% vs. ctyper 96.5%; full-set: Locityper 99.5% vs. ctyper 99.98%). If only run on 28 *HLA* genes, using one thread, ctyper takes 35 seconds (1.1s for genotyping) on genotyping all *HLA*s. As a comparison, the Locityper genotyping of *HLA* genes required about 8.5 minutes using 8 threads (4 minutes spent genotyping). Separating out the genotyping runtime from read preprocessing (which may be amortized across multiple loci), ctyper gives a 218× speedup in genotyping.

Finally, we used ctyper to genotype and repeat the previous assembly comparisons on 273 challenging medically relevant (CMR) genes^52^, of which 212 are not reported as CNV. Unrepetitive (unmasked) regions had on average 0.29 mismatches per 10 kb against the full-set; these mismatches are 99.7% less than the total variants when aligning the assembly to corresponding GRCh38 sequences (reference-based divergence). The genotypes using leave-one-out databases had 4.9 mismatches per 10 kb, with 94.8% fewer than reference variants (Supplementary Figures 4-6). Including repeat-masked low-complexity sequences (e.g., variable-number tandem repeats), there were 10.5 mismatches per 10 kb against the full-set, which is 97.6% fewer than reference-based divergence, and 74.7 mismatches per 10 kb among leave-one-out, which is 82.7% fewer than reference-based divergence (Supplementary Figures 7-9). Compared with

Locityper, ctyper works on more CMR genes (273 vs. 259) and has a lower average mismatch rate in the full-set (10.5 vs. 13.2 mismatches per 10 kb), but has a higher average mismatch rate in the leave-one-out analysis (74.7 vs. 45.4 mismatches per 10 kb). When limiting comparisons to genomic regions in common with Locityper, ctyper has improved performance (3.0 mismatches in the full-set and 22.3 mismatches in the leave-one-out analysis per 10 kb; Supplementary Table 7 and Supplementary Figure 10).

### Sequence level diversity of CNVs in global populations

We used principal component analysis (PCA) to examine the population structure of PA genotypes on 2,504 unrelated 1kGP samples, 879 Genotype-Tissue Expression (GTEx) samples, and 105 diploid assemblies (excluding HGSVC due to lower coverage after filtration) (Fig. 4a,b). Following standard population analysis, rare subgroups (<0.05 allele frequency) were excluded, and copy numbers were limited to 10 to balance the weights of principal components (PCs). The 1kGP, GTEx, and genome assemblies were clustered by population as opposed to the data source, suggesting little bias between genotyping and assembly, or across NGS cohorts.

**Figure 4.**
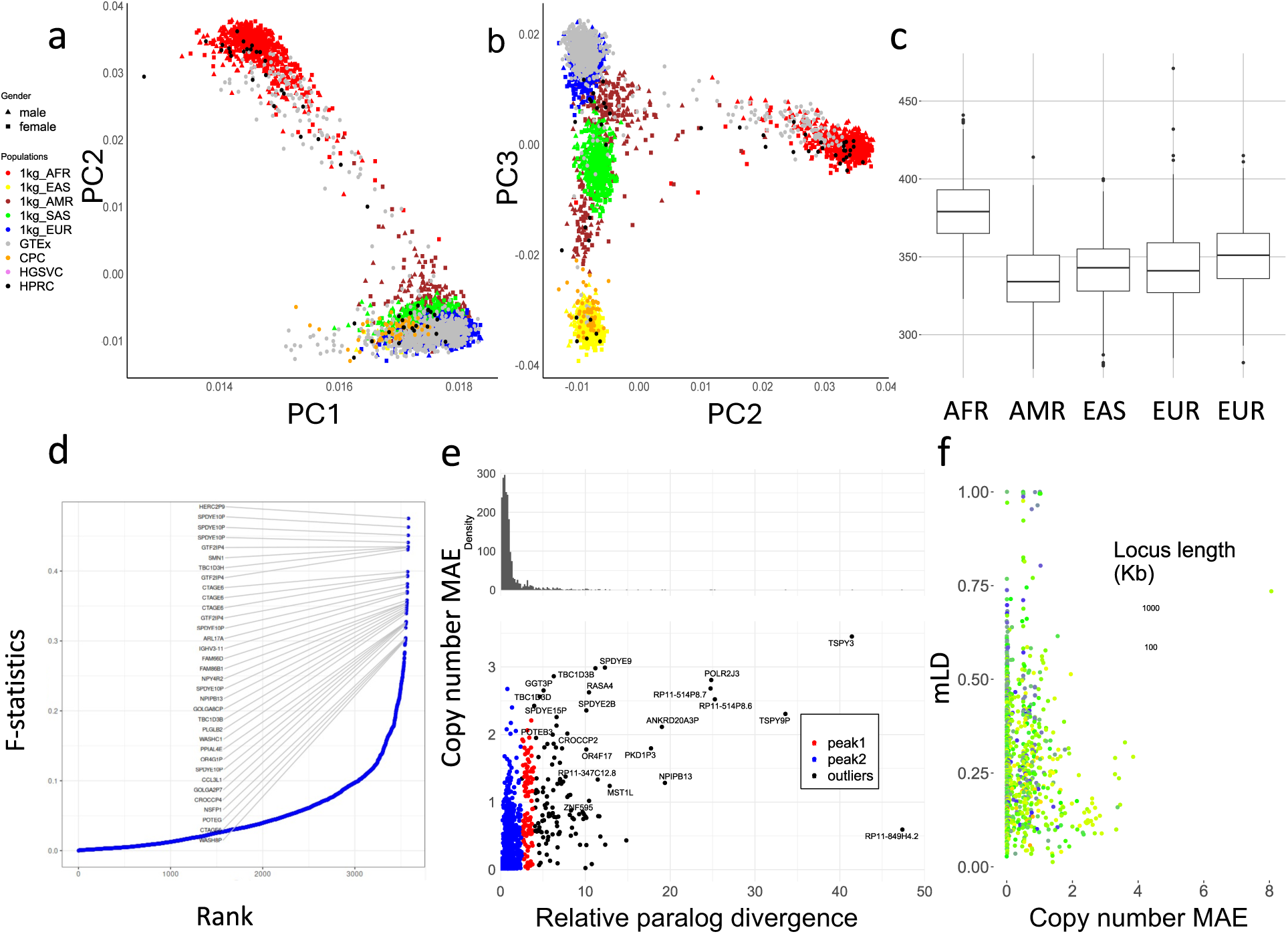
**a,b**. PCAs of allele-specific copy numbers on the union of PA genotyping results and assembly annotations. **c,** Distribution of total autosomal gene copy numbers among unrelated 1kGP samples, including pseudogenes. **d,** Population differentiation measured by F-statistics of duplications among different continental populations. The genes with a paralogous subgroup with an F-statistic of more than 0.3 are labeled. **e,** Mean absolute variation in copy numbers and relative paralog divergence in sequences. Based on our genotyping results on unrelated 1kGP genomes, for genes found to be CNV to the population median in more than 20 samples, we determined the average aggregate copy number difference (mean absolute error [MAE]) between individuals and estimated the average paralog difference in sequences relative to ortholog difference. **f,** Multi-allelic linkage disequilibrium (mLD) between pairs of CNV genes less than 100kb apart. The larger MAE value of each pair is used for the x-axis values. The total locus length denotes the length from the beginning of the first gene to the end of the last gene.

The top 0.1% highest weighted subgroups on PC1 have an average aggreCN variance of 26.33, significantly larger compared to an overall of 4.00 (p-value=1.11e-16, F-test). Similarly, PC2 and PC3 have mean aggreCN variance of 19.73 and 7.20, suggesting CNVs are weakly associated with sequence variants. Furthermore, PC1 is the only PC that clustered all samples into the same sign with a geographic center away from 0, suggesting it corresponds to modulus variance (hence aggreCN) if treating samples as vectors of paCNs. Meanwhile, PC2 and PC3 were similar to the PCA plots based on SNP data on global samples^53^, suggesting they are associated with the sequence diversity of CNV genes. The total number of duplications is elevated in African populations (Fig. 4c), reflected in the order of PC1 (Fig 4a).

We examined ctyper genotypes to measure the extent to which duplications show population specificity. We used the F-statistic, a generalization of the F_st_ that accommodates more than two genotypes (Methods), to test the differences in distributions across five continental populations (Fig. 4d). In total, 4.4% (223/5,065) of duplicated subgroups showed population specificity (F-statistic > 0.2; Supplementary Table 8). The subgroups of PAs with the highest F-statistic (0.48) contain duplications of *HERC2P9* which is known to have population differentiation^9,54^. Another example is a converted copy of *SMN2* annotated as a duplication of *SMN1* that is enriched in African populations (F-statistic=0.43).

We then measured whether duplicated genes were similar or diverged from reference copies, indicating recent or ancient duplications and providing a measure of reference bias from missing paralogs. We constructed multiple sequence alignments (MSAs; Methods) for sequences of each matrix and measured the pairwise differences in non-repetitive sequences for each pair of PAs. We determined the average paralog sequence divergence relative to ortholog divergence (Methods), which we refer to as relative paralog divergence (RPD). We also measured copy number diversity using the mean absolute error (MAE) of genes in the populations, indicating the CNV level among populations (Fig. 4e). Based on RPD, using Density-Based Spatial Clustering of Applications with Noise^55^, we identified two peaks at 0.71 and 3.2, with MAE centers at 0.18 and 0.93. The first peak indicates genes with rare and recent CNVs, while the second peak indicates more divergent and common CNVs, often CNVs that may be inherited as different structural haplotypes that cannot be analyzed using a single reference genome. For example, *AMY1A* has a high RPD at 3.10 because of the truncated duplications of *AMY1A* (blue gene annotations in Fig. 2a). If we consider the sequence variation indicates duplication age, these results are consistent with ancient bursts of duplications in humans and primate ancestors^56^.

We next studied the haplotype linkage of PAs to investigate the levels of recombination at different CNV loci. We determined multiallelic linkage disequilibrium^57^ (mLD; Methods) between PAs using the unrelated 1kGP samples for 989 subgroups that were adjacent and less than 100 kb apart on GRCh38 (Fig. 4f), and reported the average mLD within each matrix. There was a strong negative rank correlation between MAE of copy number and mLD (*ρ*=-0.24, p-value=3.4e-15, Spearman’s rank), which was stronger than the rank correlation between mLDs and total locus length (*ρ*=-0.21, p-value = 1.5e-11, Spearman’s rank), suggesting a reduced haplotype linkage on genes with frequent CNVs. The lowest mLD=0.013 was found on *FAM90*, a gene with frequent duplications and rearrangements^58^. Not surprisingly, the 29 loci with highest mLD (mLD > 0.7) are enriched in the sex chromosomes (N=19). Furthermore, *HLA-B* and *HLA-DRB* had mLD >0.7 and only deletion CNV (Supplementary Methods). In agreement with our prior diversity analysis based on long-read data, the amylase locus has a value of 0.293 due to recombination.

### Expression quantitative trait loci (eQTLs) on pangenome-derived alleles

To investigate the impact of paCNs on expression, we performed eQTL analysis using the Genetic European Variation in Disease^59^ (GEUVADIS) and GTEx^60^ cohorts. There were 4,512 genes that could be uniquely mapped in RNA-seq alignments. An additional 44 genes, such as *SMN1/2* and *AMY1A/1B/1C*, were analyzed by pooling among all copies because they have indistinguishable transcription products. We assigned PAs to these transcripts based on exonic sequences (Methods; Supplementary Table 9).

We corrected expression bias using PEER^61^ with the first three PCs from SNP genotypes^62^ and performed association analyses with paCNs. After merging paCNs to aggreCNs, 5.5% (178/3,224) of transcripts showed significance (corrected *P* = 1.6e-0.5, Pearson correlation) as previously observed^37^. We then tested whether using paCNs would provide a stronger fit by updating the aggreCNs with individual paCNs and performing multivariable linear regression on expression (Methods). There were significant improvements in fit for 27.6% (890/3,224) of transcripts (corrected *P* = 1.6e-05, one-tailed F-test; Fig. 5a).

**Figure 5.**
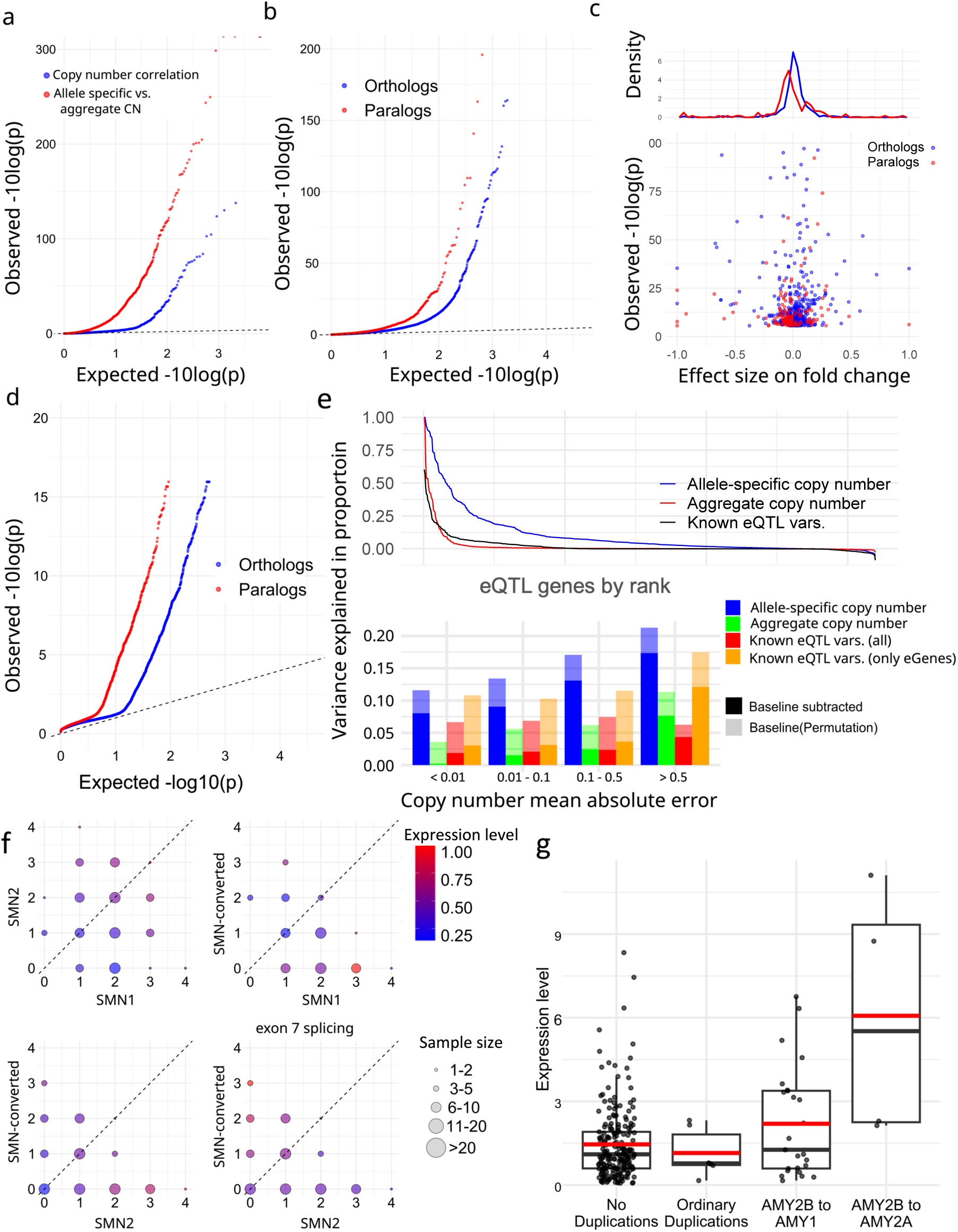
**a**. Q-Q plot of associations of (blue) aggregate copy numbers to gene expression in GEUVADIS samples (red) and the improvement of allele-specific copy number over aggreCN. **b,** Comparative gene expression Q-Q plots of orthologs (blue) and paralogs (red). **c,** Fold change effect size of all significant alternative expressions in b. The fold changes as well as p-values are shown. **d,** Preferential tissue expression of orthologs and paralogs. **e,** (top) Comparison of different models on explained expression variance (R^2). (bottom) Quantification of variance explained by different representations at different levels of CNV frequencies: full paCN genotypes, aggregate copy number, and known eQTL variants. **f,** Case study on *SMN* genes showing decreased gene expression on *SMN*-converted. The average expression level in PEER-corrected GEUVADIS samples is shown under different copy numbers of *SMN1*, *SMN2*, and *SMN*-converted. Transcript levels are the total coverage of all isoforms, and the exon 7 splicing level is measured by counting isoforms with a valid exon 7 splicing junction. **g,** Case study on amylase genes showing increased gene expression on translocated *AMY2B* using PEER-corrected GTEx pancreas data.

The improved fit could be explained by the nonuniform expression of different alleles of the same reference gene. To test this, we used a linear mixed model (LMM; Methods)^63,64^ to regress total expression to individual subgroups and estimate allele-specific expression, then compared these values to other subgroups of the same matrix that were assigned to the same reference gene (Supplementary Table 10). For subgroups within solvable matrices with >10 samples, we found that 7.94% (150/1,890) of paralogs and 3.28% (546/16,628) of orthologs had significantly different expression levels (corrected with sample size = number of paralogs + orthologs, corrected *P* = 2.7e-06, chi-squared test, Fig. 5b). Overall, paralogs were found to have reduced expression (Fig. 5c), consistent with previous findings on duplicated genes^65^.

The granularity of PA genotypes enabled testing for preferential expression of paralogs among different tissues. After correcting raw transcript per million values using DESeq2^66^, we compared expression in 57 tissues in the GTEx samples using LMMs to estimate the expression levels in each tissue (Methods; Supplementary Table 11). There was alternative tissue specificity for 132 of 2,820 paralogs (4.68%) and 225 of 19,197 orthologs (1.17%) (corrected *P* = 6.4e-08, union of two chi-squared tests; Methods; Fig. 5d).

Additionally, we used analysis of variance (ANOVA) to estimate the proportion of expression variance (R^2) explained by paCNs using PEER-corrected GEUVADIS, and compared it to a model based on known SNPs, indels, and eQTL structural variants^67^ (Methods). As expected, the highly granular paCNs explained the most variance: on average, 10.3% (14.3% including baseline). In contrast, 58.0% of transcripts are genes with known eQTL variants that explained valid variance by 2.14% (1.60% considering experimental noise, in agreement with a previous estimate of 1.97%^68^). On average, 1.98% of the variance was explained by aggreCNs, and 8.58% by subgroup information. When combining both paCNs and known eQTL sites, 10.4% (19.0% including baseline) of the valid variance was explained (Fig. 5e).

We examined the *SMN* and *AMY2B* genes as case studies due to their importance in disease and evolution^33,69^. The *SMN* genes were classified into three categories: *SMN1*, *SMN2*, and *SMN-converted*. We estimated the total expression of all transcripts and the expression of only isoforms with valid exon 7 splicing junctions. For total expression, no significant difference was found between *SMN1* and *SMN2* (0.281 ± 0.008 vs. 0.309 ± 0.009; *P* = 0.078, chi-squared test). However, significant differences were found between *SMN-converted* and *SMN1/2* (0.226 ± 0.012 vs. 0.294 ± 0.002; *P* = 1.75e-07, chi-squared test), with a 23.0% reduction in expression of *SMN-converted*. In contrast, despite having lower overall expression, *SMN-converted* had 5.93× the expression of *SMN2* (*P* = 2.2e-16, chi-squared test) for valid exon 7 splicing^70^, indicating that while *SMN-converted* has full functional splicing^71^, its overall expression level is lower (Fig. 5f).

We studied the expression of *AMY2B* duplications when they are translocated proximally to other *AMY* genes, such as the PAs containing *AMY1* and *AMY2B* in Fig. 2a. Using PEER-corrected GTEx pancreas data, we estimated their expression as well as for other duplications. We found that these translocated *AMY2B* genes had significantly higher expression than other duplications (1.384 ± 0.233 vs. −0.275 ± 0.183, *P* = 7.87e-09, chi-squared test) (Fig. 5g).

## Discussion

New pangenomes present both opportunities and challenges for the study of complex genetic variation (e.g., CNVs, recurrent SVs, translocations and gene conversions): while they reveal the landscape of complex variation, such variation requires new models for representation and analysis. We represent genomic variation as PAs: haplotype segments that capture genomic structural information and phased variation. To support large NGS cohort analyses, we developed an alignment-free genotyping tool, ctyper, to genotype PAs among NGS data, providing allele-specific sequence information as well as copy number state. The genotyping approach is based on a new mathematical model that relaxes an NP-hard problem into a more efficient polynomial semi-analytic solution while maintaining accuracy, with increased robustness and copy number sensitivity. It is also important to note that although the analysis in this study focuses on CNV genes, ctyper can also genotype non-CNV genes and provide a calling for complex genetic variation and local phasing.

The use of ctyper genotypes increases the scope of NGS studies to profile variation in challenging, medically relevant and CNV genes that are otherwise unmappable. For example, our finding that CNVs reflect two modes of variation– highly similar (and likely recent) and low-identity (ancient and polymorphic) duplications– is based on the 1kGP genotypes rather than assembly annotation. As another example, the ctyper genotypes yield tissue-specific expression of paralogs as well as relative contributions to the expression of different forms of duplications, such as *SMN*.

We investigated the significant improvement of ANOVA on PAs, whose genotypes are multiallelic and reflect different combinations of variants in contrast to known biallelic eQTL variants from conventional analysis. First, compared to PAs, there were either very few or very many eQTL variants per gene, indicating LD (Supplementary Figure 11) as addressed by fine-mapping^72^, and increasing multiple testing burden^73^. Indirect association due to LD also explains why there was a greater proportion of variance explained among genes with more CNVs by conventional eQTL variants (for example, the *HPR* genes, Supplementary Figure 2). However, as the frequency of CNVs increases, the explained variance by eQTL variants increases (t= 3.80, p-value = 1.6e-04, Pearson’s correlation), and the number of eQTL variants decreases (t = −4.79, p-value = 2.1e-06, Pearson’s correlation), suggesting that larger effects like CNVs might overshadow the discovery of other eQTL variants not in LD (the increase of total variance reduces significance in association analysis using a Gaussian-like model). Because PAs already incorporate LD information, they will suffer less from such LD-based problems. Furthermore, gene expression might not be a linear additive effect of all variants^74^. For example, although *SMN*-converted contains variants that are either from *SMN1* or *SMN2*, its overall expression is lower than both. In this manner, using a genetic model with linked variants such as PAs would improve upon the linear additive model in predicting gene expression. Because those limitations also apply to non-CNV genes, the concept of PAs may have a wider potential for future genome-wide association analysis.

Our analysis has some limitations. Due to the limited sample size, our associations are based on subgroups rather than individual PAs. Different cohort sizes may require different levels of granularity when defining subgroups. Our current classification of subgroups was designed for biobank cohorts; association tests in smaller cohorts may need to aggregate more PAs into subgroups. For example, the three subtypes of *SMN-converted* showed little difference in our eQTL analysis, so they were merged in our case study, but larger cohort studies may find their differences. The granularity of genotyping is additionally defined by the length of PA sequences; genotypes using shorter PAs will more accurately reflect small variants, while longer sequences can preserve more structural information and may be preferable in regions with low recombination, such as *HLA-DRB*.

Ctyper has limitations. First, while it is possible to detect CNVs smaller than PA units using ctyper (Supplementary Methods), full support requires additional benchmarking data. Additionally, ctyper currently does not provide confidence values for genotypes.

## Data availability

Software: https://github.com/ChaissonLab/Ctyper.

Allele database and annotations: https://doi.org/10.5281/zenodo.13381931. Benchmarking and analysis code: https://github.com/Walfred-MA/CNVAnalyze.

## Acknowledgments

This work was supported by NHGRI R01HG011649 and NHGRI U01HG010973. W.M. conceived the method, performed the analysis, and wrote the manuscript. M.J.P.C. conceived the method and wrote the manuscript. We thank Dr. Matthew Pennell for the constrictive critiques of our manuscript.

## Notes

https://zenodo.org/records/13381931

### Competing Interest Statement

The authors have declared no competing interest.

### Summary of Updates

This revision adds leave-one-out benchmarking as well as clarifications to the text.

