## Supplementary Methods for "Genotyping sequence-resolved copy number variation using pangenomes reveals paralog-specific global diversity and expression divergence of duplicated genes"

### Online Methods

#### Constructing pangenome-derived allele database

##### Search and extraction of initial genes of interest from pangenome assemblies

Our pangenome cohort was composed of assemblies from the Human Pangenome Reference Consortium (HPRC; 92 haplotypes, excluding HG02080 due to abundant flagged regions), the Chinese-Pangenome Consortium (CPC; 114 haplotypes), the Human Genome Structural Variation Consortium (HGSVC; 18 haplotypes, only PacBio HiFi assemblies were used), two telomere-to-telomere diploid assemblies (4 haplotypes), and reference genomes (GRCh38 including alternative loci and T2T-CHM13). The gene coordinates used were GENCODE v39 based on the GRCh38 reference genome.

We constructed databases for 3,203 genes found to have copy number variation in the HPRC and CPC studies. Genes were initially organized into "query sets" where each query set encompassed genes with functional or similar sequences including pseudogenes and genes with distant homology within the same gene family. The query sets were initially defined based on genes with shared name prefixes, and were used to locate all similar sequences within the pangenome.

Efficient mapping methods[1,2] missed alignments to sequences that contain *k*-mer matches that decrease genotyping accuracy when not included in our database, including small pseudogenes and diverged paralogs. To address this, we developed a sensitive and efficient scanning scheme centered on *k*-mer clusters to detect all similar sequences for genes of interest in the pangenome. For each gene or query set, we used low-copy *k*-mers (*k* = 31) derived from all initial reference genes, and that appeared fewer than 255 times in the T2T-CHM13v1.1 genome, to help locate similar sequences. We searched for these *k*-mers in each of the pangenome assemblies and references. We then identified *k*-mer hotspots defined as maximal intervals of mapped *k*-mers containing more than 200 *k*-mers within any 1,000-base window within the interval. To aid in mapping small and fragmented pseudogenes, we included an additional criterion to define hotspots: the presence of 50 exonic *k*-mers within the same interval search. Subsequently, we used BLASTn[3] to refine the boundaries of each hotspot by aligning all reference genes in this query set extended by 5,000 bases upstream and downstream flanking sequences to each *k*-mer hotspot.

For copy number variant (CNV) genes, the hotspots defined by *k*-mers often include loci mapped by multiple genes from a query set as well as tandemly duplicated genes. To account for this redundancy, we merged alignments that were less than 10,000 bases apart (together with 5,000 upstream and downstream flanking sequences- this merges genes within a distance of 20,000 bases), causing tandemly duplicated genes to be merged into a single locus. To avoid genotyping longer loci that may be split by recombination, we divided a locus at the midpoint of an intron if the intron exceeded 20 kb. To ensure a minimum locus length so that genes are comparable, flanking sequences both upstream and downstream were adjusted to achieve a minimum total length of 15,000 bases. These methods aimed to standardize the size of each sequence to be roughly 30,000 bases, approximating the size of linkage disequilibrium (LD) blocks. The collection of all sequences mapped by a query set is referred to as initial matrix-sequences.

#### **Filtration and polishing of initial matrix-sequences and *k*-mers**

For each genome, we extracted *k*-mers found in matrix-sequences and not elsewhere in the genome. From this, we also filtered out low-complexity *k*-mers with a composition of at least two-thirds redundant *2*-mers or *3*-mers. These are associated with highly repetitive DNA and cause bias in genotyping. Additionally, we excluded *k*-mers demonstrating high (>70%) or low (<30%) GC content[4]. The matrix-sequences composed of a majority of filtered *k*-mers were removed.

Next, sequences from the non-confident regions reported by the HPRC, as well as truncated sequences from small scaffolds were removed. This required sequences to be at least 10,000 base pairs away from both ends of a scaffold, except for sequences from genes located at the telomere region of the reference (<10,000 bases).

Because the initial query sets were defined solely based on gene names and included arbitrarily grouped genes with unrelated sequences, our initial groups of matrix-sequences had sequences with low-homology but similar names. Thus we used subsequent refinements to partition unrelated sequences from our initial groups of matrix-sequences.

We separated the remaining sequences into partitions based on the number of shared *k*-mers using graph partitioning. Each sequence was represented as a node, and edges were made between node pairs sharing an excess of 500 low-copy *k*-mers, except for *NBPF* and *ANKRD* genes, for which a higher threshold of 2,000 low-copy *k*-mers was set to further reduce the sizes of partitions for computational efficiency in later analyses.

After filtration and polishing, we obtained our final lists of sequences, which we labeled pangenome-derived alleles (PAs). There were 1,408,209 PAs for 3,351 genes in total distributed across 3,307 partitions. This complete list includes any additional genes not defined as duplicated in the original set, yet have high sequence similarity. The average PA length was 33 ± 29 kb and included protein-coding genes (69%), processed pseudogenes (20%), intronic duplications (5%), and decoys (unrelated genes that share homology and improve genotyping accuracy when included; 7%). We represented each final partition as a single matrix, and the list of low-copy *k*-mers specific to each matrix that passed filtration (the "*k*-mer list") was compiled together with the sequences. We label these compiled structures as "*k*-mer matrices". Each row corresponds to a PA sequence. Each column corresponds to a distinct *k*-mer. The matrix cell values are set to the counts of the corresponding *k*-mer in the respective PA. The counts are mostly 0 or 1, but occasionally more than 1 when there are low-copy repeated sequences in the PA, or the row represents a tandemly duplicated locus.

#### Annotation of pangenome-derived alleles

##### ***K-mer* based phylogenetic tree construction**

We constructed a separate phylogenetic tree for each *k*-mer matrix for both annotation and genotyping purposes. For computational efficiency as well as consistency with our *k*-mer based genotyping and annotation, we used distances based on *k*-mers instead of multiple sequence alignments (MSAs) for construction.

The matrix structure (we use M to denote any arbitrary *k*-mer matrix) allows us to easily measure the concordance between any two sequences by their vector form, $G_i$ and $G_j$, by calculating their inner product, denoted as $<G_i * G_j>$. Consequently, the norm matrix, $N = M * M^T$, reflects the *k*-mer concordances for all sequence pairs within the matrix.

We constructed a similarity matrix, S, where $S_{i,j}$ is the cosine similarity of $G_i$ and $G_j$. The cosine similarity can be obtained by normalizing the norm matrix N according to the modules of corresponding vector pairs (approximately equal to the geometric mean of the lengths of corresponding two sequences).

Finally, we used the Unweighted Pair Group Method with Arithmetic Mean (UPGMA) algorithm on the similarity matrix S to generate the phylogenetic tree for each partition.

#### Clustering of pangenome-derived alleles into highly similar subgroups

For each group of sequences corresponding to a matrix, we used its corresponding phylogenetic tree for the annotation and classification of highly similar groups of alleles, which we term “highly similar subgroups”. The classification of highly similar subgroups is guided by two primary criteria applied across all subgroups:

Homogeneity within subgroups: A subgroup must exhibit near-identical characteristics amongst its members, which is quantified by ensuring the largest *k*-mer distance between any two members does not exceed 155 *k*-mers, which is roughly equivalent to the variation caused by five single-nucleotide polymorphisms (SNPs) or a structural variant of approximately 95 bp, such that subgroups are capable of representing most common variants in about a 30 kb range.

Distinctiveness of subgroups: Each subgroup must be distinct from its neighboring subgroups. This is measured using a *k*-mer F-statistic score, which must exceed 2 when compared with adjacent subgroups. In cases where subgroups are composed of fewer than three members, the F-statistic may not be reliable; hence, we default this score to 0 for such small subgroups, but change the cutoff of the former criteria to 155 * 3 to detect singleton rare events.

Employing a “bottom-up” recursive approach starting from leaves, we applied these criteria to all clades, aiming to identify and report the largest possible highly similar subgroups. These are later used to identify equivalent loci after genotyping.

#### Pangenome-derived allele annotation relative to the reference genome

We annotated CNV events and duplicated alleles in the pangenome assemblies relative to the GRCh38 reference genome. This requires determining the corresponding GRCh38 gene for each PA. However, this is a known challenging problem of orthology assignment[5].

First, PAs often align to multiple paralogs on GRCh38, and the orthologous gene identified by reference mapping may not be the most similar reference gene due to gene conversion and translocation (Figs. 1f and 2a). To address this problem, we designed a method to match PAs to their closest GRCh38 genes based on *k*-mer similarity. For every haplotype, we obtained all pairwise similarities between each of its PAs to each of the GRCh38 PAs. We matched PAs to their most similar GRCh38 PAs, starting from the most similar pair, until all PAs were matched or failed to match (had no reference gene with >90% similarity). Secondary redundant matches (match to reference genes that had already been matched) were annotated as duplications.

Second, the formerly failed-to-match PAs were likely alleles with large structural variants, such as insertions, deletions and local proximal duplications. We attempted to map them back to GRCh38 using their flanking sequences (100 kb on either side). Because it is challenging to assign orthology regions within large segmental duplications, we designed the liftover process to be in two steps. First, we lifted PAs to the region with the best local alignment coverage, allowing SVs to break alignments into smaller units. Next, we performed a global pairwise alignment between PAs and the lifted region to locate the best-aligned gene with the presence of local translocations and tandem duplications (Supplementary Methods).

Finally, to annotate the proximal duplications as well as identify diverged paralogs that failed to match from both prior methods, we annotated PAs using gene transcripts. We aligned all exons from the same matrix to PAs, and based on the exon order and alignment scores, determined the optimal combinations of transcripts on each PA (Supplementary Methods). The PAs containing no exons were annotated as introns and PAs containing only transcripts of other unrelated genes were annotated as decoys. Introns and decoys were usually filtered out from analysis and the remaining PAs were considered as valid alleles, including pseudogenes that have no intact protein-coding transcripts and putative protein-coding genes with intact protein-coding transcripts.

#### Classification of orthologs and paralogs in the pangenome

To classify the orthology and sequence similarity of PAs with respect to reference genes, we classified PAs into four categories, including two types of orthologs and two types of paralogs:

1. Reference alleles are alleles in the same subgroup as GRCh38 alleles, representing the alleles almost identical to the reference sequences.
2. Alternative alleles are orthologs located at the same genomic locus as the reference gene but are distinctly in different subgroups from GRCh38 alleles. This

includes alleles that have high sequence divergence or large structural variants, as observed in genes like *HPR*, *NBPF*, and *CYP2D6*.

3. Duplicated paralogs (alleles) consist of paralogs that have been duplicated to different loci from their source genes, but retain high sequence similarity to the reference alleles (>80% *k-mer* similarity). These alleles reflect large, recent segmental duplications in the genome, including similar paralogs, such as *AMY1A*, *AMY1B*, and *AMY1C*, which are still often considered the same gene despite their distinct locations.

4. Diverged paralogs (alleles) consist of paralogs duplicated to different loci from their source genes and are significantly divergent (<80% in *k*-mers). These were typically characterized by highly diverse non-reference paralogs, incomplete gene duplications, and novel processed pseudogenes. An illustrative example of diverged paralogs is found among amylase genes, where there is a translocation event between *AMY1* and *AMY2B*.

#### Justification of the representation of pangenome-derived alleles and highly similar subgroups

##### Comparison of pangenome-derived alleles to other genomic representations

We characterized the information gained by representing a genome by the copy numbers of genotyped PAs (paCNs) compared to the copy number of reference alleles. For each PA, we compared the nearest neighbor in our pangenome database as a proxy for the optimal genotyping result of samples containing that PA to its closest GRCh38 gene based on *k*-mer similarity. The nearest neighbor had 94.7% fewer differences on average compared to GRCh38 matches and 57.3% had identical nearest neighbors.

We then assessed the proportion of subgroups identifiable by *k*-mers uniquely shared by all their members, analogous to SUNKs. Only 38.8% of subgroups (with at least three members) contain such *k*-mers (Fig. 2e). For example, no SUNKs exist between *SMN1*, *SMN2*, and *SMN-converted* due to gene conversion (Fig. 1f). However, there are unique combinations of *k*-mers used by ctyper genotyping.

We investigated the extent to which PAs are linked in large haplotype structures. We found that recombination or other structural variation creates unique combinations that cannot be represented during leave-one-out analysis. Using the amylase genes as an example, 40% (90/226) of haplotypes could not be represented with remaining subgroups, particularly those with a greater number of copies than GRCh38 (45/67). When all PAs devoid of SVs were considered equally in a single large subgroup, 20% (46/226) of haplotypes remained singleton, especially those with additional copies (26/67). Furthermore, many subgroups, such as the novel PAs containing both *AMY1* and *AMY2B* in proximity, are found within different structural haplotypes (Fig. 2a). While such issues may be mitigated by a larger pangenome, genotyping at the level of PA increases the ability to identify the genetic composition of next-generation sequencing (NGS) samples at highly variable multicopy gene loci.

**<u>Justification of highly similar subgroups in representing population diversity</u>**

We characterized highly similar subgroups to justify if they can well capture sufficient population diversity. The average pairwise *k*-mer cosine similarity (under this metric, one base change roughly adds up to *k* different *k*-mers) was 98.8% within each highly similar subgroup, compared to an average 94.2% cosine similarity to their corresponding reference sequence, showing a 5.03× decrease. Between two phylogenetically neighboring subgroups having at least three members each, the between-group variance is 6.03× greater than the within-group variance, showing a strong hierarchical structure. This shows that most genetic diversity may be represented using a small number of haplotype states as both criteria suggest that more than 80% of total population variation could be represented by highly similar subgroups.

#### Genotyping NGS samples with ctyper

##### Initial solution based on linear regression

Each matrix of PAs is genotyped separately. Given an NGS sample and a *k*-mer matrix M derived from PAs, we generate a vector V for an NGS sample with the counts of each *k*-mer from the sample found in the matrix, normalized by the sequencing coverage. We seek to find a vector X that denotes the copy numbers of all PAs and minimizes the squared distance to the *k*-mer counts observed in NGS data, e.g., $\mathrm{argmin}_X$ $(\| M^T * X - V \|)$. Although it is possible to directly obtain an integer solution using mixed-integer linear programing (MILP) based on absolute (edit) distance, this is NP-hard[6] and can only be efficiently used with very few variants/*k*-mers[7,8]. This restricts the use of MILP on the pangenome. However, the relaxed non-integer solution based on squared distances has an analytic solution that can be efficiently solved. Also, compared with absolute distance, squared distance is more suitable for the normal-like noise in NGS data[9,10]. In essence, the computational problem is akin to a multivariable linear regression.

To make the solution closer to the maximum likelihood estimate, during the regression, we rescaled the weights of *k*-mers to even out their expected uncertainty. Assuming the observation of *k*-mer copy number follows a negative binomial distribution with the dispersion small enough to be distinct from Poisson[10], the expected variance is roughly proportional to the square of observation, so we chose to weight the *k*-mer by the square of the reciprocal of their observed copy number. We also applied smaller weights (adjust=0.05) on singleton *k*-mers (observed in only one PA and not observed in NGS) because they are more likely to be assembly errors.

#### Integer solution based on reversed phylogenetic rounding

Initial linear regression often yields solutions in the form of small floating-point values, where the alleles with the highest coefficients are not necessarily those closest to query genes. However, as shown by mathematical analysis (Supplementary Methods), there are strong relationships between the initial least-error solution and the true integer solutions under a phylogenetic framework:

1. Non-negative solutions: Without uncertainty in predicting *k*-mer copy numbers, the least-error solution should be non-negative. Therefore, we obtain a non-negative least-error solution (NNLS) via the Lawson-Hanson algorithm[11].
2. Total copy number estimation: The sum of the initial solutions should approximate the total number of the true integer solutions, allowing us to estimate the total gene copy number in the querying sample.

3. Phylogenetic position prediction: On a binary phylogenetic tree, the branch with a shorter vector distance to the genes in the querying sample will have a larger sum of coefficients (inversely proportional to distance). This relationship enables us to predict the phylogenetic position of each gene in the querying sample.

4. Fractality of least square error solution on phylogenetic tree: if a solution is the least squared error solution of the tree, it is also the least squared error solution within each clade, allowing the greedy method to perform on the phylogenetic tree.

5. Large database effect: In large databases, having more genes highly similar to query genes increases condition number and tends to distribute the total coefficients across them, resulting in smaller individual coefficients. However, the total sum of these coefficients increases, improving the precision of phylogenetic position prediction, and this effect does not plateau.

6. With sequencing coverage variance for NGS at ~30-fold coverage, the model precision remains. Sequenced variance is not the primary source of error.

Given the high “convergence” and fractality of the solution on the phylogenetic tree in large databases, we developed a greedy algorithm to efficiently convert non-integer solutions into integer solutions. This iterative algorithm follows a bottom-up approach, starting from the leaves and progressing toward the root. At each hierarchical level, non-integer values are rounded to the nearest integer solution that minimizes the overall residual, while any remainder is propagated to the next level. Because at each level of the hierarchy there are only two remainders from either branch of the tree, this solution is highly efficient. We label this approach as reversed phylogenetic rounding. The pseudo-code for this algorithm (naive version and optimized version) is provided in the Supplementary Methods.

#### Benchmarking of genotyping

##### Hardy-Weinberg equilibrium

Hardy-Weinberg equilibrium (HWE) analysis is used to determine whether the observed proportions of homozygous and heterozygous alleles conform to Hardy-Weinberg expectations or not. After excluding sex chromosomes and setting the maximum copy number to two, we calculate the frequency of each subgroup in the population (frequency denoted as p). Based on HWE, the expected frequency of having copy number = 0 equals (1-p)^2,  the expected frequency of having copy number = 1 equals 2p(1-p), and the expected frequency of having copy number = 2 equals p^2. We compared the observations from genotyping results to expected values and determined p-values using the chi-squared distribution. The subgroups with p-values <0.05 were reported as significant.

##### Trio analysis

For trio analysis, we quantified Mendelian violations of inheritance with respect to copy-number counts. When the copy number of a child is 0, the parents need to be 0 or 1; When the copy number of a child is 1, the parents cannot both be 0 or 2; When the copy number of a child is 2, the parents both need to be 1 or 2. When the copy number of a child is more than 2, the parents need to have a gross copy number greater or equal to this number.

#### Comparison of genotyping results to pangenome assemblies

For each of the HPRC benchmarking genomes, the accuracy of PA genotypes was measured by aligning the genotyped PA sequences to the corresponding assembly (ground truth). To do so, the assembly PAs were paired one-to-one with genotyped PAs by a greedy method. We obtained all pairwise similarities measured by *k*-mer similarity between all combinations of benchmark/PA genotype pairs. Starting from the most similar pair, we paired those alleles without replacement and iterated this until all assembly PAs were either paired or failed to be paired (e.g. has no genotyped PAs with >90% similarity, considered as copy number disagreements).

Second, the paired PAs were then aligned using the global pairwise alignment tool Stretcher[12] for masked sequences and a pairwise alignment method distributed with Locityper[13] for unmasked sequences. From the global alignments, we obtained the number of mismatched bases in the unmasked region, where the low-copy repeat *k*-mers are used in *k*-mer matrices. We also compared the performance of ctyper with Locityper on unmasked sequences. The Locityper results were obtained from tables distributed on Zenodo. Additionally, we evaluated the performance of ctyper in the regions that overlap with the Locityper target loci. We first mapped the GRCh38 coordinates used to define Locityper target loci onto each pangenome assembly using minimap2 on the corresponding GRCh38 sequence, and located the mapped results on each individual

PA. When evaluating the global pairwise alignment results between PAs, we extracted the alignments corresponding to the intersection of regions mapped by minimap2 for both the target and benchmark PAs, and determined the score for the corresponding regions using the aligner included in the Locityper distribution.

#### Classification of errors

We classified four types of errors for our benchmarking:

1. False positive: the genotyping results have an additional copy;
2. False negative: the genotyping results have a missing copy;
3. Mistyping: copy assigned to an incorrect type;
4. Out of reference: the sample has a PA that is a singleton subgroup and excluded from the genotyping database during leave-one-out.

#### Benchmarking *HLA, KIR*, and *CYP2D* genes with public nomenclatures

We benchmarked the results of *HLA, KIR* and *CYP2D* genes from the 39 HPRC samples shared with 1kgp samples from both full-set and leave-one-out analyses. First, we labeled all IPD-IMGT and CYP2D-star annotations for PAs. For *HLA* and *KIR*, we annotated using Immuannot[14], and for *CYP2D6*, we annotated using Pangu[15]. Using those annotations, we converted genotyped PAs sequences into public nomenclatures and compared these with the annotation results of the assemblies from the same samples. The benchmarking results of *HLA* were compared with T1K[16] with its default settings and the benchmarking results of *CYP2D6* were compared with Aldy[7] with its default settings.

We also benchmarked SNP calling on *CYP2D6*, and compared with Aldy ran with default settings. We took the phased results of Aldy and matched them to their corresponding original PAs. In a range of about 6 kb, where the variants could be found (first SNP reported at chr22:42126309, last SNP reported at chr22:42132374), Aldy genotyped the variants with an F1 score of 85.2%, and ctyper genotyped the variants with an F1 score of 95.7%.

We ran T1K for HLA genes with commands: “run-t1k --abnormalUnmapFlag -t 12 --preset hla-wgs --alleleDigitUnits 15 --alleleDelimiter : -c hlaidx_dna_coord.fa -f hlaidx_dna_seq.fa -b $CramFile -o $Output”, and ran T1K for KIRs with commands: “run-t1k --abnormalUnmapFlag -t 12  --preset kir-wgs --alleleDigitUnits 15 --alleleDelimiter : -c kiridx_dna_coord.fa -f kiridx_dna_seq.fa -b $CramFile -o $Output”. We ran Aldy on CYP2D6 with the command: “aldy genotype -p wgs -g CYP2D6 $CramFile --reference $HG38 --output $Output”.

#### Population analysis of pangenome-derived alleles

**Total number of duplication events from genotyping results**

Based on the ctyper genotyping results, we calculated the total number of duplication events for each 1kgp unrelated sample, excluding seven samples due to having extreme values different from the population mean by more than five standard deviations. The total number of each reference gene was measured in each genome and compared to GRCh38 chromosomes, excluding alternate haplotypes. Each duplication event was called if the genome had a copy number more than twice that of

GRCh38, excluding decoys/introns and sex chromosome genes. The total number of duplication events was reported for each genome. It is important to note that these duplications also included pseudogenes and pseudogene-like exonic fragments besides known protein-coding genes.

#### Measuring F-statistic values

Because subgroups may have copy numbers beyond two and may not be applicable to the Fixation index ($F_{st}$), we instead used the F-statistic value to measure the population specificity of subgroups. The F-statistic value is based on the F-test, where we obtained the variances of copy numbers within all continental populations (within-group variance), and used it to divide the variances of copy numbers across different populations (between-group variance).

#### Relative paralog divergence

Relative paralog divergence measures the mean divergences of the paralogs to other alleles, in relation to the mean divergence between only orthologs. Relative paralog divergence was determined for each reference gene and based on the graphic MSAs (Supplementary Methods) of PAs assigned to that reference gene as well as the ctyper genotyping results.

First, the divergence value was determined for each pair of PAs assigned to the same reference gene. It was measured based on the alignment scores of unmasked bases (misalignment and gap open = -4, and gap extend = 0, normalized by total alignment length) from graphic MSAs.

Second, we obtained the mean divergence of the orthologs by averaging divergence values between the two PAs from samples with CN = 2.

Third, we then determined the population median copy numbers for each reference gene, and divided samples into those with additional copy numbers (copy numbers more than the median) and those with no additional copy numbers (copy numbers not more than the median).

It may be unreliable to directly distinguish the paralog from orthologs due to complex rearrangements (e.g. Fig. 2a). To overcome this limitation and only obtain the divergence values from additional copies, we performed statistical estimation based on large populations. We first estimated the mean divergence values from samples with no additional copy numbers and used this as the unit baseline B. When the population median CN = Y, because there are Y(Y-1)/2 pairs, then the total baseline is B * Y(Y-1)/2, which will be subtracted from total divergence values of samples with duplications, and Y(Y-1)/2 will be subtracted from the total number of pairs (the denominator) as well. For a sample with CN = X, the estimated paralog divergence is [ Total_Variance - B * Y(Y-1)/2 ] / [ X(X-1)/2 - Y(Y-1)/2 ].

After subtracting the total baseline, the mean paralog divergence value of the additional copies was determined for all samples with additional copy numbers. The mean paralog divergence was then normalized by the mean divergence of the orthologs obtained in step two.

##### Multi-allelic linkage disequilibrium

Multi-allelic linkage disequilibrium (mLD) is an analytic continuation of SNP-based bi-allelic LD to allow computing linkages between multiple genotypes on neighboring loci. When there are only two genotypes on both loci, mLDs equal the LD value. When there are more than two genotypes, we measure LD between each pair of genotypes across both loci and take the weighted average of all pairs. This weight is the product of both allele frequencies of the pair.

#### Expression analysis of pangenome-derived alleles

##### Determining transcripts for expression analysis

We first represented each gene by their major transcripts from the MANE (Matched Annotation from NCBI and EMBL-EBI[17]) project. Second, individual exons were aligned. Transcripts were recursively clustered together if they overlapped with previously clustered transcripts with more than 98% overall similarity taking the average similarity of all aligned exons from the transcripts. We considered these clusters as the same transcript even though they were from different genes. Third, for each transcript, we identified all its exons and looked for unique exons that did not overlap with exons from other transcripts. Fourth, we used these unique exons to represent each transcript and filtered out transcripts that had no unique exons (2,079 out of 2,579 filtered genes were known pseudogenes). Last, we assigned PAs to each transcript if they contained any of the corresponding unique exons with at least 98% similarity.

#### Expression correction

For individual tissue analysis, similar to the prior study[18], we logistically corrected the raw transcript-per million (TPM) values using the tool PEER together with the first three principal components. For Geuvadis samples, we obtained PCs from PLINKv2[19] with default settings on the reported genotypes in chr1[20], and for GTEx samples, we obtained PCs from the GTEx database. For cross-tissue analysis, we corrected raw TPMs using DESeq2[21] with default settings.

#### Association between CNVs to gene expression

We first associated gene aggregate copy number to expression levels using Pearson correlation (linear fitting). The p-values and residuals of this fit were recorded. To test if including allele-specific information would improve the fitting, we used the ctyper PA-specific copy numbers to replace the aggregate copy numbers to perform multi-variable linear regression using PA-specific copy numbers as dependent variables, and gene expression level as independent variables. We compared the residuals of multi-variable linear regression with residuals from the former linear fitting using the F-test. The one-tailed p-values of the reduced residual were reported. Both p-values were corrected by the number of transcripts tested (N=3,224).

#### Linear mixed model

We performed linear mixed modeling (LMM) to measure the individual expression of each subgroup. We used the total gene expression values as the vector of observed dependent variables, different subgroups as the vector of independent fixed variables, and the copy numbers from the ctyper genotyping results were used as their coefficient matrix. The effect sizes of fixed variables were then solved using ordinary least squares (OLS) regression.

#### Alternative expression of subgroups

We determined whether a subgroup has an alternative expression level compared to other subgroups of the same gene. Paralogs were assigned to a reference gene based on exonic annotation, where "gene" explicitly refers to those in GRCh38. To avoid redundancy with previous studies using a single reference genome, in this section, we did not compare expressions between known reference paralogs, such as *SMN1* and *SMN2*. We merged all other subgroups assigned to the same reference gene into a single variable separate from the subgroup currently being tested. Additionally, we included other factors, such as paralogs assigned to other reference genes that might also influence total expression, as additional parameters to adjust for their potential interference. For subgroups within solvable matrices with more than ten non-zero expressions, using an LMM and the R lm function[22], we regressed the expression values to all variables to get their effect sizes. We then compared the effect size of the currently tested subgroup and the effect size of other subgroups of the same gene using the chi-squared distribution with the linearHypothesis package[23]. This p-value was then corrected by the number of total subgroups tested (N=18,518).

#### Across-tissue expression comparison

We determined if a subgroup has an alternative most-expressed tissue compared to other subgroups of the same gene. Paralogs were assigned to a reference gene based on exonic annotation, where “gene” explicitly refers to those in GRCh38. To avoid redundancy with previous studies using a single reference genome, in this section, we did not compare expressions between known reference paralogs, such as *SMN1* and *SMN2.* we merged all other subgroups assigned to the same reference gene into a single variable, separate from the currently tested subgroup. Additionally, we included other factors, such as paralogs assigned to other reference genes that might also influence total expression, as additional parameters to adjust for their potential interference. For subgroups within solvable matrices with more than 10 non-zero expression values, we used a linear mixed model to estimate the gene expression level of each subgroup within each of the 57 tissues in GTEx V8. The tissue with the highest expression level was recorded and compared to the tissue with the second highest expression using the chi-squared test. We then compared the results between the currently tested subgroup and all other subgroups of the same gene to see if they had a different highest-expressed tissue. When the highest expressed tissues were different, we tested the p-value of either event by combining the p-values from both sides as p-combined = p1 + p2 - p1 * p2. This p-value was then corrected by the number of subgroups tested on all 57 tissues (N=776,902).

#### ANOVA (Analysis Of Variance) tests on gene expression

We first measured the total expression variance for each expression quantitative trait locus (eQTL) transcript, filtering out units with per-sample variance less than 0.1 to exclude genes not sufficiently expressed in the Geuvadis cohort. We estimated experimental noise by measuring expression variance between different trials of the same individuals (mean = 10.5% of the total variance) and excluded transcripts with experimental noise exceeding 70% of the total variance, resulting in 639 total transcripts after filtration. We applied the one-in-ten rule to restrict the number of variants tested to be not greater than 45 (10% of the sample size) to avoid over-fitting. We filtered out 18 transcripts involving more than 45 PAs. When there were more than 45 known eQTL variants, we used 45 variants with the lowest p-values. The valid expression variance was obtained by subtracting experimental noise from the total expression variance. Using ANOVA, we estimated the explained valid variance and adjusted the results by subtracting a baseline, defined as the mean expression variance explained by permuting the orders of all samples (estimated by the mean of 100 trials). If there were no reported eQTL variants, a value of 0 was used for known eQTL variants.

We further investigated the part of variance explained by gene aggregate copy-number (aggreCN). We applied ANOVA to random matrices with aggreCN information, with randomly assigned subgroups that have the total copy number equal to the original matrix. We then subtracted the variance explained by these random matrices (estimated by the mean of 100 trials) from the total explained variance by paCNs to obtain the variance explained by subgroup information.

1. Li, H. Minimap2: pairwise alignment for nucleotide sequences. *Bioinformatics* **34**, 3094–3100 (2018).
2. Ren, J. & Chaisson, M. J. P. lra: A long read aligner for sequences and contigs. *PLoS Comput. Biol.* **17**, e1009078 (2021).
3. Altschul, S. F., Gish, W., Miller, W., Myers, E. W. & Lipman, D. J. Basic local alignment search tool. *J. Mol. Biol.* **215**, 403–410 (1990).
4. Ross, M. G. *et al.* Characterizing and measuring bias in sequence data. *Genome Biol.* **14**, R51 (2013).
5. Kirilenko, B. M. *et al.* Integrating gene annotation with orthology inference at scale. *Science* **380**, eabn3107 (2023).
6. Hartmanis, J. Computers and intractability: A guide to the theory of NP-completeness (Michael R. garey and David S. johnson). *SIAM Rev. Soc. Ind. Appl. Math.* **24**, 90–91 (1982).
7. Numanagić, I. *et al.* Allelic decomposition and exact genotyping of highly polymorphic and structurally variant genes. *Nat. Commun.* **9**, 828 (2018).
8. Ford, M. K. B. *et al.* ImmunoTyper-SR: A computational approach for genotyping immunoglobulin heavy chain variable genes using short-read data. *Cell Syst* **13**, 808–816.e5 (2022).
9. Hodson, T. O. Root-mean-square error (RMSE) or mean absolute error (MAE): when to use them or not. *Geosci. Model Dev.* **15**, 5481–5487 (2022).
10. Daley, T. & Smith, A. D. Predicting the molecular complexity of sequencing libraries. *Nat. Methods* **10**, 325–327 (2013).
11. Lawson, C. L. & Hanson, R. J. Back Matter. *Solving Least Squares Problems* 312–

337 Preprint at https://doi.org/10.1137/1.9781611971217.bm (1995).

12. Rice, P., Longden, I. & Bleasby, A. EMBOSS: the European Molecular Biology Open Software Suite. *Trends Genet.* **16**, 276–277 (2000).

13. Prodanov, T. *et al.* Locityper: targeted genotyping of complex polymorphic genes. *bioRxiv* (2024) doi:10.1101/2024.05.03.592358.

14. Zhou, Y., Song, L. & Li, H. Full resolution HLA and KIR gene annotations for human genome assemblies. *Genome Res.* (2024) doi:10.1101/gr.278985.124.

15. GitHub - PacificBiosciences/pangu. *GitHub* https://github.com/PacificBiosciences/pangu.

21. Love, M. I., Huber, W. & Anders, S. Moderated estimation of fold change and

dispersion for RNA-seq data with DESeq2. *Genome Biol.* **15**, 550 (2014).

22. Fox, J. & Weisberg, S. Mixed-effects models in R. *An R Companion to Applied Regression; SAGE: Thousand Oaks, CA, USA* (2002).
23. Fox, J., Weisberg, S. & Price, B. Car: Companion to applied regression. *CRAN: Contributed Packages* The R Foundation https://doi.org/10.32614/cran.package.car (2001).
